# Indirect versus Direct Effects of Freshwater Browning on Larval Fish Foraging

**DOI:** 10.1101/804070

**Authors:** Dina M. Leech, Troy L. Clift, Jessica L. Littlefield, Nicholas R. Ravagli, Jacob E. Spain

**Affiliations:** Department of Biological and Environmental Sciences, Longwood University Farmville, VA 23901

**Keywords:** brownification, fish larvae, foraging, predation, zooplankton

## Abstract

Fish foraging and energy flow are both predicted to decline with freshwater ‘browning’ due to reductions in light availability. Studies investigating these predictions have focused on juveniles and adults; however, the larval stage represents a ‘critical period’ in fish development. We investigated the indirect versus direct effects of browning on zooplankton-larval fish interactions by altering water color with SuperHume (absorbance at 440 nm = 1.6 – 10.8 m^−1^). Phytoplankton and zooplankton densities were monitored across experimental tanks in the laboratory for one month leading up to fish spawning. Larval largemouth bass were then introduced to assess indirect effects on fish feeding rates and growth. Direct effects on foraging of largemouth bass and bluegill were determined with separate short-term feeding experiments. Browning did not directly alter the ability of larval fish to capture prey. However, significant indirect effects on larval fish foraging, growth, and survival were observed as phytoplankton and zooplankton decreased with increased browning. Our data suggest lake browning will reduce energy transfer to larval fish due to a reduction in prey availability but not visual foraging.

## Introduction

In recent decades, many freshwater and coastal ecosystems in the Northern Hemisphere have become browner in color due to an increased export of chromophoric dissolved organic matter (cDOM) from the terrestrial watershed (Monteith et al. 2007; Erlandsson et al. 2008; Haaland et al. 2010; Solomon et al. 2015). The mechanisms underlying this ‘browning’ are currently debated and include changes in climate, hydrology, and land use, reduced atmospheric acid deposition, and increased inputs of dissolved iron (Freeman et al. 2001; Monteith et al. 2007; Erlandsson et al. 2008; Kritzberg and Ekström 2012). Whatever the mechanism, this phenomenon has far reaching ecological consequences for the structure and function of aquatic ecosystems, including energy flow.

One concern is the effect of browning on the underwater light environment. As waters become browner in color, the quantity of light in the water column is reduced, resulting in a shallower euphotic zone (Bukaveckas and Robbins-Forbes 2000; Einem and Graneli 2010). In addition, shorter wavelength ultraviolet and visible radiation are more readily absorbed by cDOM compared to longer wavelengths (Morris et al. 1995; Wetzel 2001). Consequently, the light environment shifts to the red portion of the visible spectrum. These alterations in the quantity and quality of light have the potential to affect zooplankton-fish interactions--indirectly through reduced energy transfer up the food chain and directly through reduced foraging efficiency.

While recent studies have shown that the initial browning of low productive systems can stimulate primary production (Ask et al. 2012; Seekell et al. 2015; Williamson et al. 2015), excessive browning of fresh waters can reduce rates of photosynthesis due to competition for photons (Kirk 1994). cDOM has been reported to sequester 10 times more photons within the visible spectrum (400 - 700 nm) compared to photosynthetic pigments (Thrane et al. 2014). As a consequence, areal primary production often decreases with increased water color (Jones 1992; Carpenter et al. 1998; Ask et al. 2009; Karlsson et al. 2009; Thrane et al. 2014), and pelagic primary production typically exceeds benthic primary production (Vasconcelos et al. 2018; Vasconcelos et al. 2019). Phytoplankton community composition can also shift to predominantly cyanobacteria, which are generally less nutritious or inedible to zooplankton (Ekvall et al. 2013; Robidoux et al. 2015). Overall, aquatic ecosystems often become net heterotrophic with increased browning as bacterial production exceeds primary production (Cole et al. 1994; Ask et al. 2012).

Reductions in the quantity and quality of basal resources then ‘cascade up’ the grazer food chain to influence zooplankton and fish. For example, Robidoux et al. (2015) observed decreases in crustacean zooplankton biomass and density with increased water color while Craig et al. (2017) found that bluegill in lakes of increasing color were smaller in size and had lower fecundity compared to those living in lakes with less color. Taipale et al. (2018) noted that zooplankton and fish had poorer nutritional quality in browner systems because the phytoplankton on which they fed had lower concentrations of essential fatty acids, proteins, lipids, and carbohydrates. Combined, these results suggest that freshwater browning indirectly alters energy flow to higher trophic levels.

Trophodynamics between larval fish and zooplankton are particularly important as the larval stage represents a critical phase in fish development that ultimately affects population growth and biomass (Fuiman and Werner 2002; Karlsson et al. 2009; Karlsson et al. 2015). Generally, in temperate climates, increased primary production in the late spring/early summer leads to increased zooplankton production. Shortly thereafter, many fish species begin to spawn, matching larval fish hatching with an abundance of zooplankton prey (Mills et al. 1989; Mehner and Thiel 1999; Hansson et al. 2007). However, as fresh waters continue to brown and energy flow to zooplankton is reduced, fish larvae may compete for fewer zooplankton prey.

Freshwater browning may also directly affect fish foraging behavior by altering the light environment to which fish are adapted. Many fish are visually orienting predators, depending on the quantity (i.e. intensity) and quality (i.e. spectra) of underwater light to locate and capture prey (Guthrie and Muntz 1993; Leech and Johnsen 2009). Results from studies investigating the direct effects of browning on fish foraging are varied, ranging from no effect to enhanced effects (e.g. Stasko et al. 2012; Jönsson et al. 2013; Weidel et al. 2017). However, this research has focused on juvenile and adult life history stages, with no knowledge of the effect of browning on fish larvae.

Foraging in early life history stages may be particularly affected by freshwater browning due the rapid attenuation of shorter wavelength light. Over evolutionary time, the visual system of fishes spectrally tunes to the intensity and wavelengths of light present in their environment, varying with age and behavior (Douglas and Djamgoz 1990). Larval fish spend most of their time foraging in the top few meters of the epilimnion, and many species have been shown to possess UV photoreceptors during only their early stages of development (reviewed in Leech and Johnsen 2009). Thus, fish larvae may rely on short-wavelength ultraviolet and blue radiation to forage (Leech and Johnsen 2009 and references therein). Alternatively, as light levels decline, fish larvae may rely on other sensory mechanisms to forage, such as olfactory cues or mechanoreception, as demonstrated in marine fish larvae and zebrafish (Jones and Janssen 1992; Cobcroft and Pankhurst, 2003; Sampson et al. 2013; Carillo and McHenry 2016).

Here, we use laboratory experiments to assess the indirect versus direct effects of browning on larval fish foraging at the time of hatching (i.e., late spring/early summer in central Virginia, USA). Based on the primary literature, we hypothesized that increased browning will reduce phytoplankton biomass, leading to a reduction in zooplankton abundance, and consequently larval fish foraging efficiency, growth, and survival during an early, critical stage in development. We then used the same laboratory set up to examine the direct effects of browning on fish feeding rate and prey selectivity when given equal zooplankton prey, hypothesizing that fish larvae will consume less prey as water color increases due to a reduction in light availability for foraging.

## Methods

### Indirect Effects of Browning

Laboratory experiments were conducted in twelve 20 L glass aquaria assembled on three shelves with four tanks per shelf. The entire shelving unit, including individual shelves, was covered in black plastic with additional black plastic placed in between each tank. Experimental tanks were arranged in a randomized block design, with at least one tank from each treatment on each shelf. Nine tanks were filled with 17 L of artificial lake water made from the COMBO medium recipe, which provides macro- and micronutrients in relatively high concentrations to support growth (Kilham et al. 1998). SuperHume, a commercially available source of humic acid, was then added to the tanks at varying concentrations to adjust the brown color of the water. Three tanks received 7 μg/L of SuperHume to serve as a light brown treatment (water color measured as absorbance at 440 nm (a_440_) = 1.6 m^−1^), three tanks received 33 μg/L to serve as a moderate brown treatment (a_440_ = 5.7 m^−1^), and three tanks received 66 μg/L to serve as a dark brown treatment (a_440_ = 10.8 m^−1^). While SuperHume adds carbon to the system, it does not add nutrients. Thus, despite differences in water color, and consequently light transparency, our treatments had similar nutrient concentrations. We adjusted the pH of all tanks to ^~^7 with approximately 2 μL of 2N HCl.

Although the experimental use of SuperHume has been previous tested (Lennon et al. 2013; Övergaard 2019), some have cited toxicity concerns for zooplankton (Robidoux et al. 2015). We therefore filled the remaining three tanks with 17 L of lake water from Sandy River Reservoir, Farmville, VA (a_440_ = 1.5 m^−1^) to serve as a comparison with our light brown SuperHume treatment. Lake water was passed through a GF/F filter to remove bacterio-, phyto-, and zooplankton, as best as possible. Sandy River Reservoir water was not used as the base water for all browning treatments because of the logistics of transporting and filtering the necessary large volume of water needed for the experiment.

Light was provided by 1.2 m long grow lamps containing two Lumichrome ^®^ Full Spectrum Plus fluorescent, 32W bulbs suspended approximately 10 cm above the water surface of the tanks on each shelf and placed on a 14-hr light, 10-hr dark cycle. While lamps cannot exactly reproduce the solar spectrum, these specific bulbs were selected based on their broad coverage of the ultraviolet and visible spectra. An Ocean Optics Red Tide USB650 UV spectrophotometer was used to measure the light spectrum, based on photon counts, from 250 to 800 nm in each treatment (Figure 1). After the experiment had concluded, we acquired a LiCor LI-192 quantum sensor to estimate photosynthetically active radiation (PAR) across our treatments. The sensor was placed in the center of the tank at a depth of 0.09 m. PAR ranged from approximately 15 μmol m^−2^ s^−1^ in the dark brown treatment, 22 μmol m^−2^ s^−1^ in the moderate brown treatment, and 31 – 35 μmol m^−2^ s^−1^ in the lake water and light brown treatments. These light levels are comparable to those experienced during crepuscular periods, when larvae often actively forage (Keast and Welsh 1968; Leech and Johnsen 2009). They are also representative of light levels experienced at midday in the summer between approximately 1.0 – 2.5 m depth in local, natural systems (i.e. ^~^ 1800 μmol m^−2^ s^−1^ surface irradiance, Figure S1). All tanks stabilized to a room temperature of 22 - 23 °C under both light and dark conditions.

**Figure 1.**
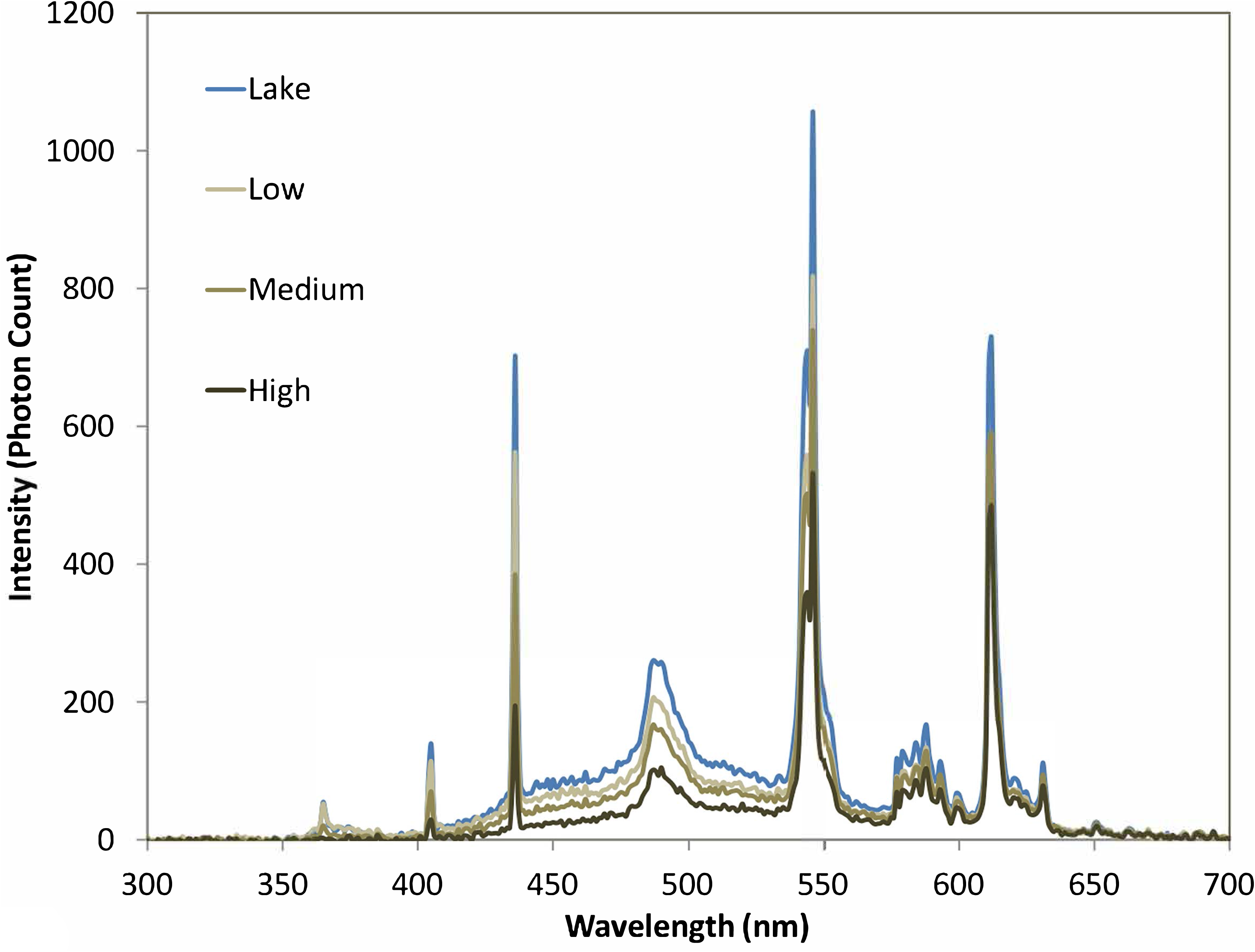
Spectral characteristics of the four treatments measured in photon counts per wavelength. Data were collected by an Ocean Optics Red Tide USB650 UV spectrophotometer from 250 to 800 nm with the sensor placed at the bottom of the tank pointing straight up toward the overhead grow lamps.

At the beginning of the experiment, each tank received a dense mixture of phytoplankton containing equal parts of *Ankistrodesmus* sp., *Chlorella* sp., *Scenedesmus* sp., and *Selenastrum* sp., which resulted in an initial chlorophyll-*a* concentration of approximately 10 μg/L in all tanks. Algae were purchased from Carolina Biological and cultured in COMBO medium prior to the experiment. A mixed assemblage of zooplankton was then added to each tank. Zooplankton were collected from Sandy River Reservoir the day before the experiment by towing a 64 μm mesh bongo net from 0 - 6 m several times. Prior to introduction, zooplankton were concentrated into a single 4 L container, mixed, and then 50 mL aliquots were introduced into each tank to provide a starting density of approximately 26 zooplankton per liter in each tank. This initial relatively low density permitted observations of population growth over time and reflected natural zooplankton concentrations in the early spring in local systems. Proportionally, the zooplankton assemblage consisted of 80% copepods (mostly cyclopoids), 15% cladocerans (*Daphnia* sp. and *Bosmina* sp.), and 5% rotifers (*Keratella* sp.). We removed all *Chaoborus* before introducing the zooplankton to the tanks to minimize zooplankton mortality due to predation.

Over the course of the next month, phytoplankton biomass and zooplankton abundance were measured at 0, 3, 7, 14, 21 and 28 days. Before each sampling, tanks were gently mixed with a broad, plastic spoon. For phytoplankton, two replicate 50 mL water samples were removed from each tank and filtered through Whatman GF/F filters to measure chlorophyll-*a* concentration as a proxy for phytoplankton biomass. Filters were placed in 90% acetone overnight in the freezer and then chlorophyll-*a* concentration was measured on a Shimadzu Trilogy Fluorometer using the non-acidification module. Chlorophyll-*a* concentrations are reported as the average of these two replicates. One hundred milliliters of DI water were added to each tank to replace the 100 mL removed for chlorophyll-*a* analysis, keeping the volume at ^~^17 L.

For zooplankton, 3 L of water from each tank was passed through a 64 μm mesh cup, and then the water was immediately returned to the tank to maintain a constant volume. Zooplankton collected on the mesh were rinsed into a sample cup and preserved with 70% ethanol. Zooplankton were identified and enumerated under a dissecting microscope at 40x in a Ward counting wheel. Zooplankton density (individuals per liter) was determined by counting the total number of rotifers, copepods, and cladocerans in each sample collection and then dividing by the total volume of water sampled (i.e. 3 L).

Because bacteria can serve as an alternate food source for zooplankton, either directly or through the microbial loop (Sanders and Porter 1990; Wylie and Currie 1991), we measured bacterial abundance at Days 0 and 28. Five milliliters of water were collected from each tank with a serological pipet, placed in a sterile culture tube, and preserved with glutaraldehyde. Samples were refrigerated until analysis. Bacterial abundance was determined by counting DAPI stained cells under an epifluorescent microscope at a magnification of 100x, based on the methods of Porter and Feig (1980). A minimum of 400 cells were counted per sample to determine cell density per milliliter.

After 28 days, two larval largemouth bass (*Micropterus salmonoides*), 10-12 mm in standard length (i.e. body length excluding tail), were introduced to each tank. Preliminary experiments determined the fish behaved better in pairs compared to single introductions. We chose to introduce the larvae after one month to simulate the timing of fish hatching in nature following an increase in algae and zooplankton in late spring (Mills et al. 1989; Mehner and Thiel 1999; Hansson et al. 2007). Because we were unable to successfully obtain fish larvae from Sandy River Reservoir prior to the experiment, we used larvae from a local pond with similar optical characteristics. Fish larvae were collected using light traps (Aquatic Instruments, Inc., Hope, ID) anchored in the littoral zone of the pond overnight. Fish were immediately transported back to the lab and housed for 24 hours without food. Fish collection and care followed approved institutional animal care and use protocols.

Once introduced to the experimental tanks, the fish larvae were allowed to feed for 24 hours. After which, zooplankton were sampled as described above. This provided an estimate of daily zooplankton consumption in each treatment, assuming that total zooplankton consumed was represented by the difference in zooplankton density at the beginning and end of fish feeding. We also assumed that the two fish in each tank fed equally, such that the difference in zooplankton density was divided by 2 to estimate daily consumption rates per fish larva.

Fish were then allowed to feed in the tanks for another 5 days with survival monitored daily. After which, fish larvae were euthanized in MS-222, and final measurements were taken of their body length under a dissecting microscope to estimate growth. Because we could not clearly distinguish the two fish in each tank, growth was estimated as the average size of the two fish at the beginning versus the end of the experiment. A final collection was also made of zooplankton density in the tanks. However, zooplankton abundance was too low in all the tanks to make accurate counts.

Throughout the experiment, a YSI 600 XLM sonde was used to measure water temperature, conductivity, pH, and dissolved oxygen concentration in each tank. To monitor potential changes in water color over time, a Shimadzu UV/Vis spectrophotometer was used to measure absorbance at 440 nm of water filtered through a GF/F filter. Dissolved organic carbon (DOC) concentration was also monitored during the experiment using GF/F filtrate. Samples were run on a Shimadzu TOC-L analyzer.

### Direct Effects of Browning

The same experimental setup was used to investigate direct effects of browning on fish foraging efficiency. However, we did not use a lake water control. Light to dark brown water color treatments consisted of four replicate tanks assembled in a randomized, block design on the three shelves. After adding SuperHume to the tanks, we noted that water color ranged from a_440_ = 1.6 to 13.1 m^−1^. In each tank, the pH was adjusted to approximately 7, and the water was allowed to equilibrate to room temperature (22 - 23°C). A YSI 600 XLM sonde was used to determine the water temperature, conductivity, dissolved oxygen concentration, and pH of each tank prior to the beginning of each feeding experiment.

Both zooplankton and larval *Micropterus salmonoides* were collected from Sandy River Reservoir. Fish larvae were starved for 24 hours prior to the experiment. For each experimental tank, two larval fish (^~^13 mm in length) were introduced and allowed to acclimate for a minimum of 1 hour. After which, a known concentration of zooplankton prey (i.e. ^~^ 20 zooplankton/L) was added, and the fish were allowed to feed for 30 minutes. We staggered the introduction of the zooplankton every 10-15 minutes to allow time for disassembling each tank at the end of the timed feeding trial. The zooplankton community consisted of approximately 40% cladocearans (mostly *Daphnia* sp.), 27% adult calanoid copepods, 24% *Chaoborus*, 7% copepodids, and 2% adult cyclopoid copepods.

Fish were removed from the tank and placed in MS-222 for euthanization. The remaining zooplankton were collected by filtering the water in each tank through a 64 μm mesh. Zooplankton were rinsed off the mesh and into a sample cup with 70% ethanol. Zooplankton were identified and enumerated under a dissecting microscope at 10 - 40x in a Ward counting wheel. Zooplankton density per liter was calculated as the total number of counted zooplankton divided by the total volume of water in the tank (i.e. 17 L). The total zooplankton consumed per minute by each fish larvae was then calculated as the difference in zooplankton density at the beginning versus the end of an experiment divided by 30 minutes and then divided by 2 larvae. We assumed each fish in each tank fed equally during the experiment. We chose this method rather than examining gut contents to be consistent with previous experiments. Preliminary studies filling and emptying tanks with zooplankton resulted in ^~^99% recovery, providing confidence in our methodology (unpublished data).

To further explore the direct effects of browning on larval fish foraging, we performed three additional experiments using the same experimental setup but with bluegill *Lepomis machrochirus* (^~^13-15 mm in standard length) collected from a local pond. The goal of these experiments was to test the limits of larval fish foraging under increasing water color by reducing the foraging time and number of prey. For each experiment, fish larvae were collected with light traps 24 - 48 hours before the experiment. Two larvae were placed in each tank and allowed to acclimate for 24 hours prior to the introduction of zooplankton prey. A YSI sonde was used to confirm that temperature, pH, conductivity, dissolved oxygen concentrations were similar across all tanks. For the first experiment, fish larvae fed for 24-hours with a relatively high density of prey (i.e., 130 zooplankton per liter) to estimate daily feeding rates under ideal conditions. We then conducted two experiments with reduced time and prey concentration to test the limits of larval fish foraging: 1) a 10-minute feeding trial with 4 zooplankton per liter and 2) a 5-minute feeding trial with 2 zooplankton per liter. Zooplankton prey remaining in each tank were collected at the end of each experiment to assess fish feeding rates and prey selectivity. For all three experiments, zooplankton were collected from the same pond as the fish larvae and consisted of approximately 80% cladocearans (mostly *Daphnia* sp.), 15% cyclopoid copepods, 3% *Chaoborus*, and 2% rotifers (*Keratella* sp. and *Asplanchna* sp.).

### Statistical Analyses

All statistical tests were performed using the R Statistical Environment (R Core Team 2018). For the indirect effects experiment, we used the *nlme* package (Pinheiro et al. 2020) to conduct repeated measures ANOVAs in combination with post hoc Tukey tests to assess differences in water temperature, conductivity, pH, and dissolved oxygen with browning treatment, time, and the interaction between time and treatment. Temperature and conductivity data were log transformed for normality. ANOVAs were computed as linear mixed models using the *lme* function, including terms for random effects associated with tank number (i.e. *random* function) and, in some cases, autoregressive effects associated with time points being unequally spaced (i.e. *corAR1* function). The *anova* function from the *car* package was used to report the results of the models (Fox and Weisberg 2019). If significant, we used the *cld* function in the *lsmeans* package (Lenth 2016) to summarize the Tukey results. Residuals from each test were calculated using the *residuals* function and then plotted with the *plotNormalHistorgram* function from the *rcompanion* package (Mangiafico 2020). Akaike Information Criteria were used to select the best model, particularly if autoregressive correlations improved the model.

Repeated measures ANOVA with post hoc Tukey tests were also performed on log transformed chlorophyll-*a* concentrations to assess differences in algal biomass with treatment, time, and interactions between treatment and time. The test was performed as described above. In addition, linear regressions of chlorophyll-*a* concentrations in each treatment over the first week (Days 0, 3, and 7) were performed to determine initial phytoplankton growth rates, using the slope as an estimate of added algal biomass per day.

For the zooplankton, we observed similar densities across treatments except on the last sampling date. We therefore assessed differences in zooplankton density only on Day 28 with a Welch’s ANOVA. This test is recommended when the data have high heteroscedasticity and is paired with a post hoc Games-Howell test to compute pairwise comparisons of treatments. We used the *welch_anova_test* function from the *rstatix* package (Kassambara 2020). Results from the Welch’s ANOVAs were used to compute effect sizes based on omega squared values.

A two-way ANOVA combined with a post hoc Tukey test was used to compare bacterial abundance across treatments between Day 0 and Day 28. A Shapiro-Wilk’s test confirmed data normality prior to running the ANOVA. The *lme* function in the *nlme* package was used to run the model, using tank number as a random variable. Summary results were presented with the *anova* function and residuals were checked with the *residuals* and *plotNormalHistogram* functions. Results of the Tukey test were observed using the *lsmeans* and *cld* functions as described above.

For the fish data in the indirect effects experiment, we again observed unequal variance across treatments, and therefore, used Welch’s ANOVAs with post hoc Game-Howell tests to assess potential differences in zooplankton consumption and growth with water color. Because we noted variability in zooplankton abundance within our treatments, we also calculated Pearson correlation coefficients between zooplankton prey availability at the time of fish introduction and zooplankton consumption rates after 24 hours as well as larval fish growth after 6 days. E*Indices were calculated to determine potential differences in prey selectivity with browning (Lechowicz 1982). Values were checked for normality and then a two-way ANOVA was performed to test for significant differences in prey selectivity between treatment, zooplankton taxa, and interactions between treatment and taxa. Tukey pairwise comparisons were performed for significant results.

For the direct effect experiment with largemouth bass, we observed differences in the absorbance coefficients within treatments. We therefore used linear regression analysis to assess the relationship between zooplankton consumption and water color (i.e. absorbance at 440 nm). For the bluegill experiments, absorbance coefficients were similar within treatments, and thus, we assessed differences in zooplankton consumption across water color treatments with Welch’s ANOVAs, using omega squared values to estimate effect sizes. E*Indices were again calculated, and two-way ANOVAs were performed to assess significant differences in prey selectivity with treatment, zooplankton taxa, and their interaction. For the largemouth bass experiment, this required us to bin tanks into light, moderate, and dark brown treatments based on similarities in a_440_, with 4 replicates per treatment.

## Results

### Indirect Effects of Browning

Over the course of the 28-day experiment, there were no significant differences in water temperature, pH, or dissolved oxygen concentration among the 12 tanks with time or treatment (Table 1). Conductivity did not significantly vary with time but was significantly lower in the lake water treatment compared to the SuperHume treatments (Table 1). Absorbance at 440 nm was similar in the light brown and lake water treatments but significantly differed in the moderate and dark brown treatments (Table 1). Water color decreased during the first week in all treatments and then remained relatively constant (Figure S2). On average, absorbance coefficients at 440 nm were 1.3 m^−1^ in the light brown and lake water treatments, 5.0 m^−1^ in the moderate brown treatment, and 9.7 m^−1^ in the dark brown treatment. DOC concentration significantly differed among the four treatments, with the light brown treatment having the lowest DOC concentration (2.6 mg/L) and the lake water treatment having the highest DOC concentration (5.1 mg/L) (Table 1). DOC concentration initially increased by 0.3 – 0.8 mg/L over the first week of the experiment and then declined by ^~^0.5 mg/L over the next 3 weeks, except in the lake water treatment, which did not significantly differ over the next three weeks (Figure S2). Interestingly, only marginal increases in DOC concentration were observed with SuperHume additions despite an approximate 10 times increase in water color. Higher DOC concentrations in the lake water treatment, compared to the SuperHume treatments, were due to high inputs of non-chromophoric, algal-derived organic carbon in the eutrophic reservoir.

**Table 1.** Mean ± S.E. of major physical and chemical parameters across water color treatments. Significance is denoted in the last column, with, T = treatment effect, D =Day, I = Treatment:Day interaction, and NS = not significant. Lowercase letters indicate which treatments were significantly different from one another based on treatment effect.

|  | Lake Water | Light Brown | Moderate Brown | Dark Brown | Significance |
| --- | --- | --- | --- | --- | --- |
| Temperature (°C) | 22.1 ± 0.18 | 22.09 ± 0.16 | 22.04 ± 0.16 | 22.03 ± 0.15 | (T) NS<br>(D) NS<br>(I) NS |
| Dissolved Oxygen (mg/L) | 9.05 ± 0.09 | 9.12 ± 0.09 | 8.89 ± 0.04 | 8.83 ± 0.03 | (T) NS<br>(D) <0.0001<br>(I) NS |
| Conductivity (mS/cm) | 0.08 ± 0<br>a | 0.25 ± 0.01<br>b | 0.25 ± 0<br>b | 0.25 ± 0.01<br>b | (T) <0.0001<br>(D) 0.01<br>(I) NS |
| pH | 7.36 ± 0.04 | 7.25 ± 0.04 | 7.15 ± 0.22 | 7.14 ± 0.02 | (T) NS<br>(D) NS<br>(I) NS |
| a440 | 1.77 ± 0.65<br>a | 0.99 ± 0.08<br>a | 4.84 ± 0.12<br>b | 9.83 ± 0.17<br>c | (T) <0.0001<br>(D) 0.003<br>(I) NS |
| Chl-*a* (ug/L) | 16.34 ± 2.69<br>ab | 29.65 ± 3.83<br>b | 12.4 ± 2.12<br>a | 8.66 ± 1.57<br>a | (T) <0.0001<br>(D) 0.004<br>(I) 0.0002 |
| DOC (mg/L) | 5.22 ± 0.05<br>a | 2.58 ± 0.07<br>b | 3.40 ± 0.06<br>c | 4.66 ± 0.07<br>d | (T) <0.0001<br>(D) 0.02<br>(I) NS |

Bacterial abundance significantly differed with time (F_(1)_=32.95, p=0.0004, n=12) and treatment (F_(3)_=24.11, p < 0.0001, n=12). Based on the results of the Tukey test, bacterial abundance in the lake water treatment (i.e., 2.86 × 10^6^ cells per mL ± 1.4 × 10^5^ S.E.) was significantly greater than all other treatments (i.e., approximately 7.1 × 10^5^ cells per mL ± 1.5 × 10^5^ S.E. in the light brown treatment and approximately 1.3 × 10^6^ cells per mL ± 9 × 10^5^ S.E. in the moderate and dark brown treatments). This suggests that not all bacteria were removed from the lake water treatment during the initial set-up, and bacteria were introduced with the addition of SuperHume. By the end of the experiment, bacterial abundance did not significantly differ across the light, moderate, and dark brown treatments but was significantly higher in the lake water treatment (i.e., ^~^ 3.19 × 10^6^ cells per mL in the lake water treatment compared to ^~^ 2 × 10^6^ cells per mL in the SuperHume treatments). Interestingly, bacterial abundance differed by only 10 ± 9% between Day 0 and Day 28 in the lake water treatment while in the low, moderate, and dark brown treatments, bacterial abundance was 64 ± 5%, 35 ± 5%, and 23 ± 3% higher on Day 28, respectively.

Phytoplankton biomass, as estimated by chlorophyll-*a* concentration, significantly differed with treatment (F_(3)_= 33.79, p < 0.0001, n=12), time (F_(1)_= 8.07, p=0.004, n=12), and the interaction between treatment and time (F_(3)_=20.18, p=0.0002, n=12) (Table 1; Figure 2). The highest chlorophyll-*a* concentrations throughout the experiment were observed in the light brown treatment (i.e., 46.1 μg/L by Day 28, Figure 2) while there was no significant difference in the moderate brown, dark brown and lake water treatments (i.e., declining to 2 −7 μg/L by Day 28, Figure 2). Based on slopes from regression analyses, algal growth during the first week was fastest in the light brown treatment (5.61 μg/L per day ± 0.25 S.E.) followed by the lake water (4.15 μg/L per day ± 0.22 S.E), moderate brown (2.36 μg/L per day ± 0.28 S.E), and dark brown treatments (1.51 μg/L per day ± 0.2 S.E).

**Figure 2.**
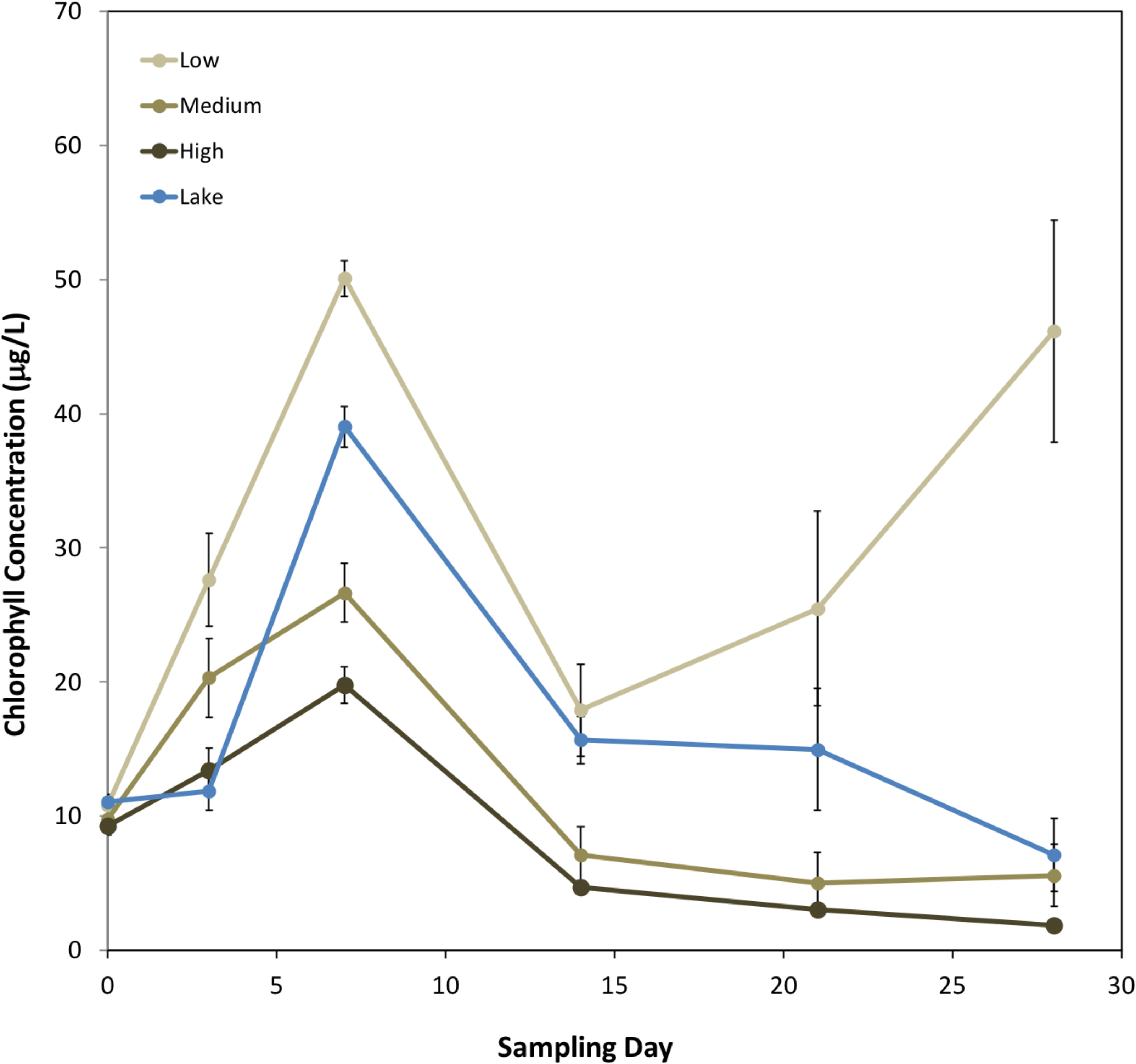
Mean phytoplankton biomass, as estimated by chlorophyll-*a* concentration, in the four treatments during the 28-day experiment. Error bars represent the standard error of three replicate tanks per treatment. Phytoplankton biomass increased in all treatments during the first 7 days but was comparatively lower in treatments with increased water color throughout the experiment.

After 28 days, total zooplankton abundance was significantly different between treatments (F_(3,3.35)_=34.4, p=0.005, n=12, est ω^2^= 0.89: 95% CI [0.0, 0.97]). Based on post hoc pairwise comparisons, zooplankton in the lake water (mean = 190 individuals per L ± 25 S.E.) and light brown (mean = 120 individuals per L ± 35 S.E.) treatments were not significantly different (p>0.05), but the lake water treatment was significantly greater than the moderate brown (mean = 72 individuals per L ± 7 S.E., p=0.02) and dark brown treatments (mean = 47 individuals per L ± 10 S.E., p = 0.003) (Figure 3A). Removing one outlier from the light treatment resulted in significant differences between all three SuperHume treatments at the p < 0.01 level. Overall, the cladoceran *Bosmina* sp. was the most abundant zooplankton in all treatments by Day 28 (Figure 3B). Despite being the dominant zooplanktors at the beginning of the experiment, copepods represented only 12% of the zooplankton community in the lake water treatment and 5% of the zooplankton community in the SuperHume treatments by Day 28 (Figure 3D & 3E). *Daphnia* sp. abundance was low in all treatments by Day 28, except for the light brown treatment (Figure 3C). While counting the zooplankton under the microscope, flocculant SuperHume was observed, but not quantified, in the guts of cladocerans (i.e., *Daphnia* and *Bosmina*) but not calanoid or cyclopoid copepods (Figure 4). Rotifers rapidly decreased in abundance and were not observed in any of the treatments after Day 7.

**Figure 3.**
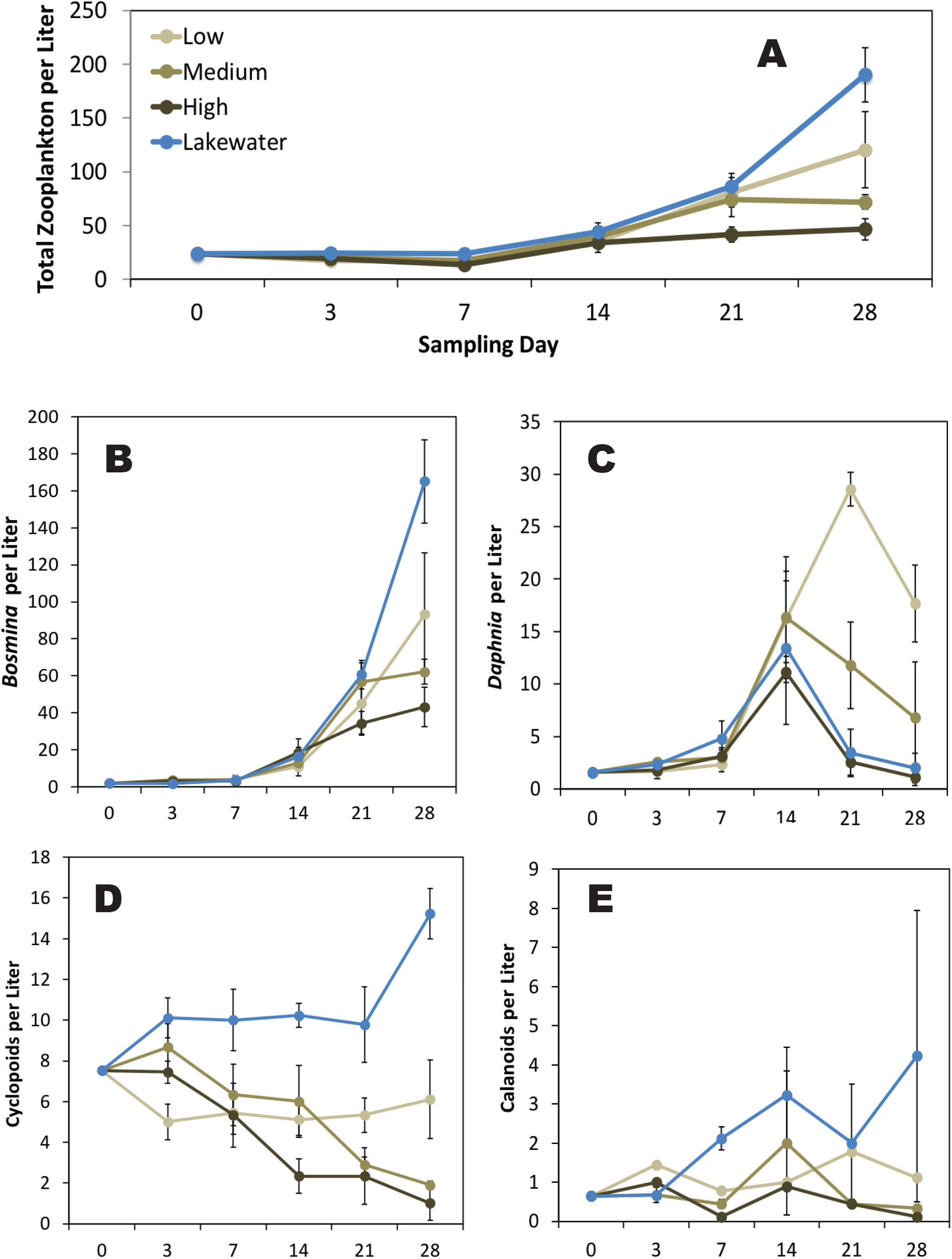
Zooplankton densities (individuals per Liter) in the four treatments during the 28-day experiment. Error bars represent the standard error of three replicate tanks per treatment. The top panel displays changes in total zooplankton densities (A) while the bottom panels display individual zooplankton genera or groups, including the cladocerans *Bosmina* (B) and *Daphnia* (C) as well as cyclopoid (D) and calanoid (E) copepods. In general, the zooplankton community shifted from copepod to cladoceran dominated and abundance was significantly lower in treatments with increased water color.

**Figure 4.**
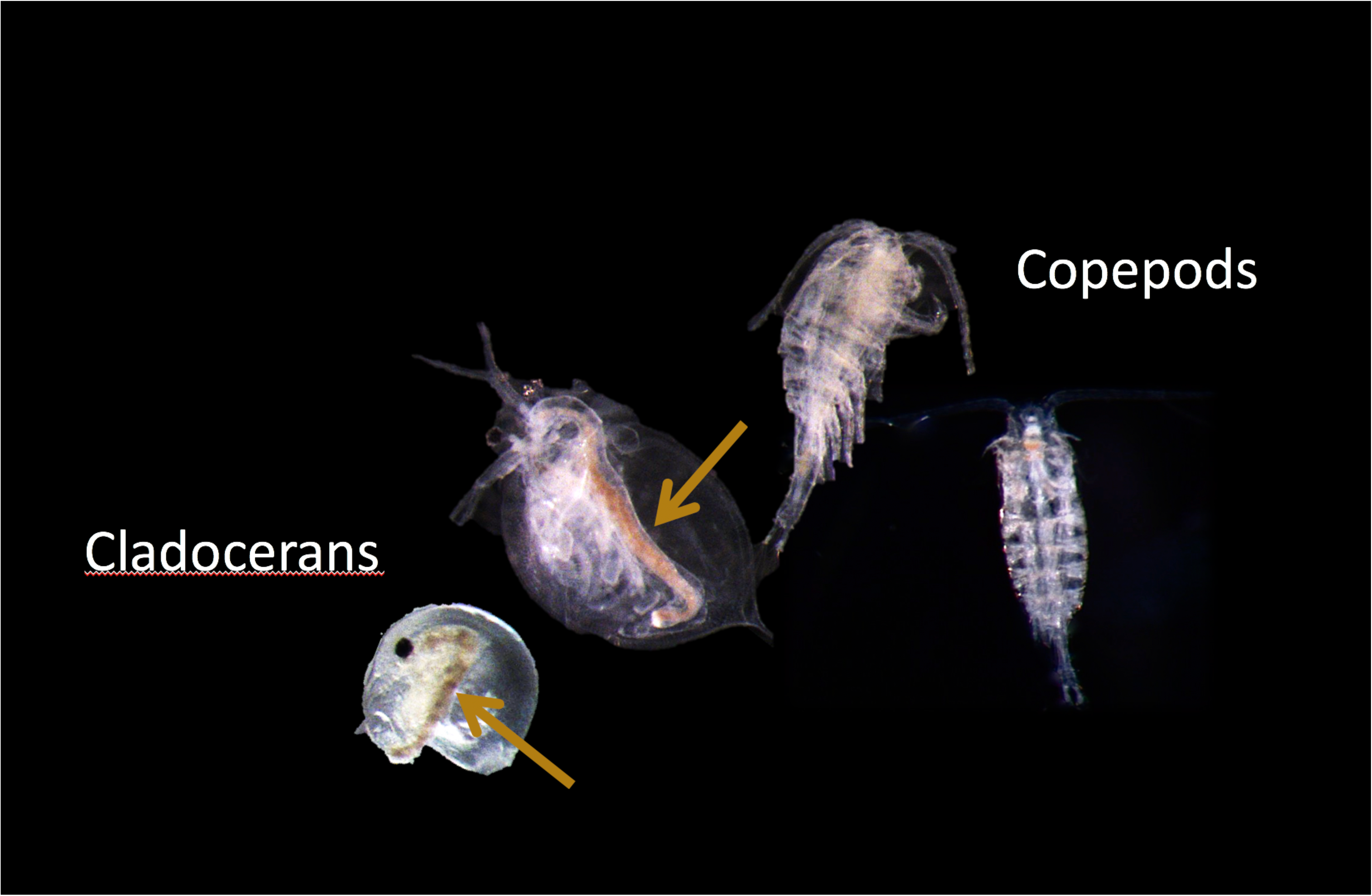
Flocculant organic matter was observed in the guts of cladocerans, including *Bosmina* and *Daphnia*, but not copepods in the moderate and dark brown treatments. Brown arrows highlight cladoceran gut contents. Image taken at 40x magnification.

All fish survived the 6-day incubation in the lake water and light brown treatments. However, there was a 33% mortality rate in both the moderate and dark brown treatments. One dead fish was found in two replicate tanks of each treatment after 5-6 days, at which point most zooplankton were visually depleted in the tanks. In the first 24 hours, fish consumed twice as many prey in the lake water treatment (i.e. ^~^610 zooplankton per larva) compared to the light brown treatment (^~^323 zooplankton per larva) and four times as many prey than in the dark brown treatment (^~^143 zooplankton per larva). However, because of the variability in zooplankton abundance within treatments, zooplankton consumption by larval fish did not significantly differ with water color (F_(3,3.49)_=4.79, p=0.09, n=12, est ω^2^= 0.48: 95% CI [0.0, 0.78]). Rather, zooplankton consumption in the first 24 hours correlated highly with zooplankton densities at the time fish were introduced to the experimental tanks (Pearson Correlation Coefficient =0.88, P=0.0002; Figure 5A). Proportionally, fish larvae primarily consumed *Bosmina* in all treatments as they were the most numerous species in the tanks. However, *Daphnia* had a higher electivity value (E*= 0.24 ± 0.0004), followed by *Bosmina* (E*= −0.12 ± 0.02) and then copepods (E*= −0.64 ± 0.02). These differences in selectivity of zooplankton taxa were statistically significant based on a two-way ANOVA (F_(2,20)_=250.9, p < 0.0001, n=12). No statistical differences in electivity were observed across treatment (F_(2,20)_= 0.44, p = 0.78, n=12) or the interaction between treatment and taxa (F_(8,20)_=0.64, p = 0.73, n =12). Note that one dark brown tank had to be eliminated from the two-way ANOVA because of a lack of *Daphnia* at the time the fish were introduced.

**Figure 5.**
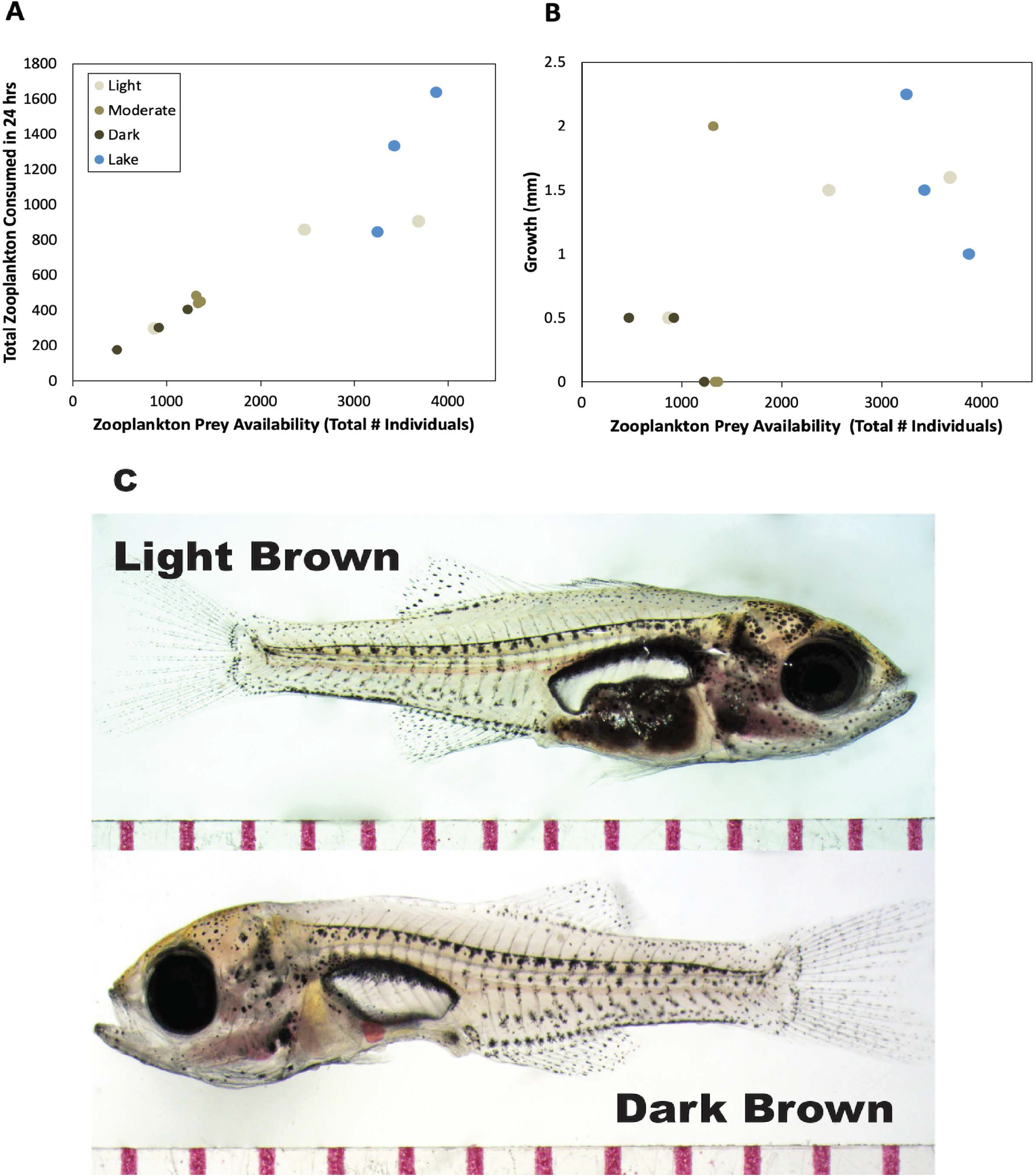
Daily prey consumption (A) and total growth over 6 days (B) in relation to zooplankton prey availability at the time fish were introduced to the experimental tanks. Note that there are 2 points for the moderate brown treatment with ^~^1400 zooplankton prey and 0 mm growth. After 6-days, surviving larval fish in the moderate and dark brown treatments displayed concave empty stomachs while those in the light brown and lake water treatments had full stomachs. Photos show example larvae from the light brown and dark brown treatments (C). Red tick marks below fish are in millimeters.

Fish growth over the 6-day feeding experiment was generally greater in the lake water and light brown treatments compared to the moderate and dark brown treatments; however, these differences were not significantly different (F_(3,4.02)_=3.17, p=0.15, n=12, est ω^2^= 0.35: 95% CI [0.0, 0.68]). On average, fish grew ^~^1.2 – 1.5 mm in the lake water and light brown treatments, ^~^0.66 mm in the moderate brown treatment, and ^~^0.33 mm in the dark brown treatment. Growth was moderately correlated with initial zooplankton abundance in the tank (Pearson Correlation Coefficient =0.62, P=0.03; Figure 5B). After 6 days, the surviving fish larvae in the moderate and dark brown treatments displayed empty and concave stomachs while those in the lake water and light brown treatments were full with zooplankton prey (Figure 5C). Note that previous studies have reported that it takes 4 – 6 hours for larvae to fully evacuate their guts (Werner 1969).

### Direct Effects of Browning

In the experiment with largemouth bass, we observed no significant differences in fish foraging efficiency with increased browning over the 30-minute feeding trial based on linear regression (Figure 6). Fish consumed on average 2.5 zooplankton per minute ± 0.3 S.E. across all treatments, consuming mostly *Daphnia* sp. (percent consumed = 61% ± 0.03 S.E.) and *Chaoborus* sp. (percent consumed = 57% ± 0.07 S.E.) followed by copepods (percent consumed = 0.07% ± 0.02 S.E.). E*Index values significantly differ across zooplankton taxa (F_(2,27)_= 71.14, p < 0.0001, n=12), with *Daphnia* sp. (E* Index= 0.19 ± 0.02 S.E.) and *Chaoborus* sp. (E* Index = 0.15 ± 0.03 S.E.) prey preferred over copepods (E* Index= −0.7 ± 0.07 S.E.). However, prey selectivity did not significantly differ with treatment (F_(2,27)_= 0.07, p = 0.93, n=12) or the interaction between treatment and zooplankton taxa (F_(4,27)_= 0.38, p = 0.82, n=12).

**Figure 6.**
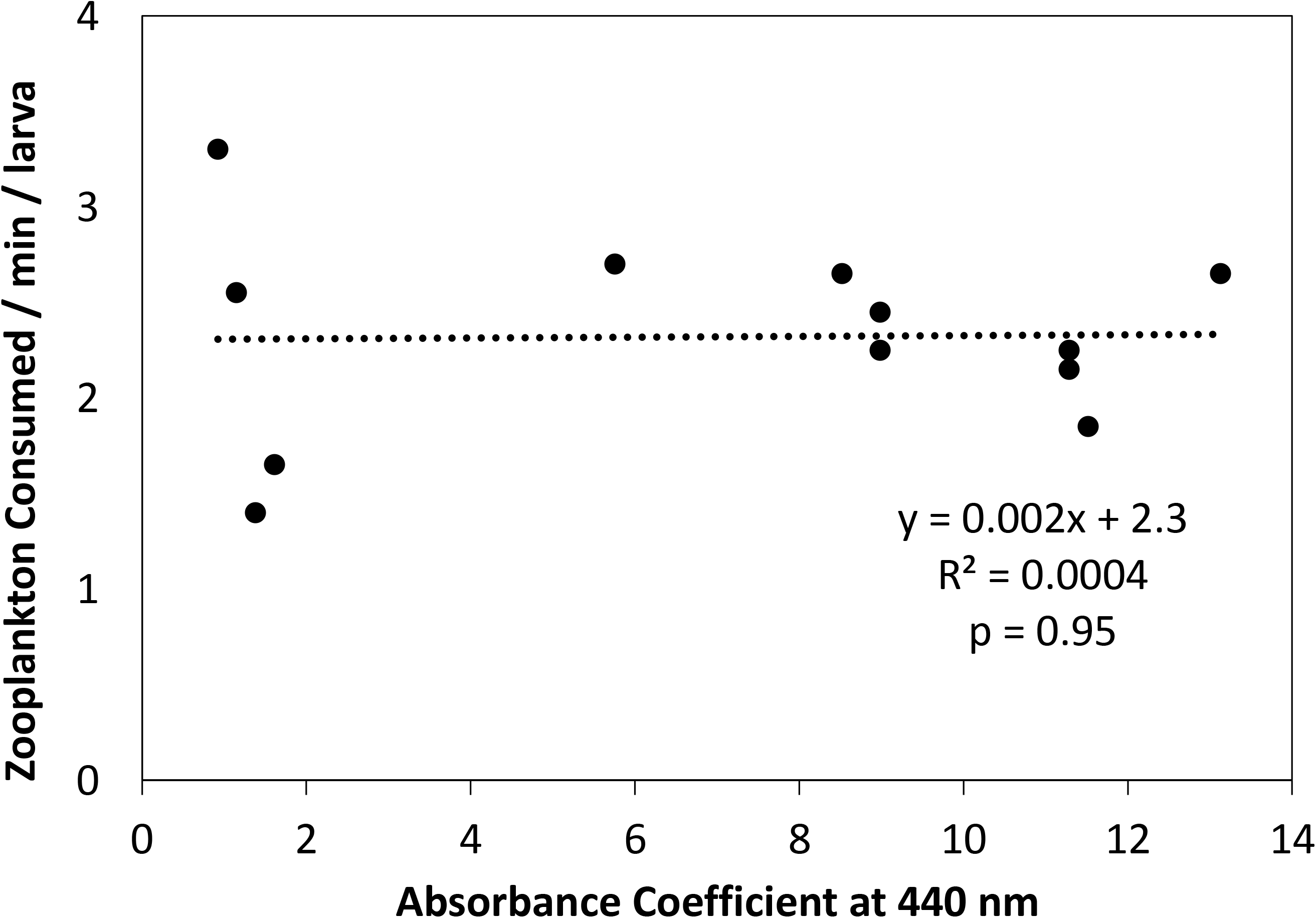
Zooplankton consumption rates for larval largemouth bass with increasing water color, as measured by absorbance at 440 nm. When given equal prey concentrations in all treatments (i.e., 20 individuals per liter), there was no significant difference in the number of prey consumed during the 30-minute feeding trial. This suggests no direct effect of browning on visual foraging.

Similar patterns were observed in the experiments with bluegill (Tables 2 and 3). Despite reducing zooplankton prey concentration and foraging time, no significant differences in foraging rate were detected with increasing water color (Table 2). Similar to largemouth bass, larval bluegill primarily consumed *Daphnia* sp. and *Chaoborus* sp. prey compared to copepods in all treatments (Table 3). In the 24-hour experiment, larval bluegill in all treatments consumed ^~^80% of the zooplankton in their respective tanks, with low consumption rates for rotifers (Table 2).

**Table 2.** Results from the three foraging experiments with larval bluegill (*Lepomis machrochirus*), including the mean ± S.E. consumption rates. The approximate total number of each zooplankton type in the tanks at the start of a feeding trial is provided in parentheses in the first column. Welch’s ANOVA results and omega-squared estimates of effect size were calculated for total zooplankton consumed only.

|  | Consumption Rates<br>(day<sup>-1</sup> larva<sup>-1</sup> or min<sup>-1</sup> larva<sup>-1</sup>) |  |  |
| --- | --- | --- | --- |
|  | Light | Moderate | Dark |
| **Experiment 1: 24 hours, ~130 zooplankton L<sup>-1</sup>** |  |  |  |
| Large Cladocerans (210) | 99 ± 4 | 101 ± 2 | 102 ± 1 |
| Small Cladocerans (1817) | 884 ± 7 | 876 ± 8 | 876 ± 11 |
| *Chaoborus* (33) | 23 ± 0 | 23 ± 0 | 23 ± 0 |
| Copepods (47) | 16 ± 0.1 | 16 ± 0.2 | 16 ± 0.2 |
| Rotifers (77) | 33 ± 1 | 34 ± 1 | 35 ± 0.1 |
| Total (2183) | 1055 ± 6 | 1051 ± 9 | 1052 ± 10 |
| Welch's ANOVA | F<sub>(2, 5.7)</sub> = 0.081, p = 0.92, n = 12 |  |  |
| Omega Squared Estimate | est ω<sup>2</sup> = 0.18: 95% CI [0.0, 0.68] |  |  |
| **Experiment 2: 10 minutes, ~4 zooplankton L<sup>-1</sup>** |  |  |  |
| Large Cladocerans (39) | 2 ± 0.03 | 2 ± 0.1 | 2 ± 0.03 |
| *Chaoborus* (5) | 0.3 ± 0 | 0.3 ± 0 | 0.25 ± 0 |
| Copepods (23) | 1 ± 0.02 | 1 ± 0.04 | 0.8 ± 0.1 |
| Total (67) | 3 ± 0.01 | 3 ± 0.2 | 2.8 ± 0.4 |
| Welch's ANOVA | F<sub>(2, 4.0)</sub> = 1.53, p = 0.32, n = 12 |  |  |
| Omega Squared Estimate | est ω<sup>2</sup> = 0.08: 95% CI [0.0, 0.47] |  |  |
| **Experiment 3: 5 minutes, ~2 zooplankton L<sup>-1</sup>** |  |  |  |
| Large Cladocerans (22) | 1.3 ± 0.2 | 1.5 ± 0.08 | 1.4 ± 0.2 |
| Small Cladoceran (7) | 0.3 ± 0.1 | 0.5 ± 0.1 | 0.38 ± 0.1 |
| Copepods (11) | 0.5 ± 0.2 | 0.7 ± 0.2 | 0.5 ± 0.1 |
| Total (40) | 2 ± 0.3 | 2 ± 0.2 | 2 ± 0.5 |
| Welch's ANOVA | F<sub>(2, 5.3)</sub> = 1.45, p = 0.31, n = 12 |  |  |
| Omega Squared Estimate | est ω<sup>2</sup> = 0.08: 95% CI [0.0, 0.43] |  |  |

**Table 3.** E* Index values from the three foraging experiments with larval bluegill (*Lepomis machrochirus*), including the mean ± S.E. for prey selectivity. The E* Index ranges from 1 to −1, where 1 represents a strong selection for a prey type, −1 equals strong avoidance of a prey type, and 0 equals random feeding. Two-way ANOVAs were performed to assess significant differences with treatment, zooplankton taxa, and their interaction.

|  | E* Index |  |  |
| --- | --- | --- | --- |
|  | Light | Moderate | Dark |
| **Experiment 1: 24 hours, ~130 zooplankton L<sup>-1</sup>** |  |  |  |
| Large Cladocerans | -0.007 ± 0.02 | 0.004 ± 0.004 | 0.007 ± 0.006 |
| Small Cladocerans | 0.01 ± 0.005 | 0.003 ± 0.005 | 0.003 ± 0.005 |
| *Chaoborus* | 0.02 ± 0.002 | 0.02 ± 0.005 | 0.02 ± 0.002 |
| Copepods | 0.01 ± 0.003 | 0.006 ± 0.003 | -0.01 ± 0.005 |
| Rotifers | -0.04 ± 0.02 | -0.04 ± 0.01 | -0.03 ± 0.004 |
| ANOVA Results: Treatment: $F_{(2,45)} = 0.09$; $p = 0.91$, $n = 12$<br>Zooplankton Taxa: $F_{(2,45)} = 20.62$; $p < 0.0001$, $n = 12$<br>Treatment:Zooplankton: Taxa: $F_{(8,45)} = 0.84$; $p = 0.57$, $n = 12$ | | | |
| **Experiment 2: 10 minutes, ~4 zooplankton L<sup>-1</sup>** |  |  |  |
| Large Cladocerans | 0.004 ± 0.008 | 0.006 ± 0.02 | -0.03 ± 0.06 |
| *Chaoborus* | 0.01 ± 0.002 | 0.04 ± 0.02 | 0.06 ± 0.06 |
| Copepods | -0.02 ± 0.009 | -0.05 ± 0.007 | -0.06 ± 0.03 |
| ANOVA Results: Treatment: $F_{(2,27)} = 0.10$; $p = 0.91$, $n = 12$<br>Zooplankton Taxa: $F_{(2,27)} = 4.81$; $p = 0.02$, $n = 12$<br>Treatment:Zooplankton: Taxa: $F_{(4,27)} = 0.72$; $p = 0.58$, $n = 12$ | | | |
| **Experiment 3: 5 minutes, ~2 zooplankton L<sup>-1</sup>** |  |  |  |
| Large Cladocerans | 0.16 ± 0.07 | -0.01 ± 0.07 | 0.10 ± 0.06 |
| Small Cladocerans | -0.23 ± 0.26 | -0.05 ± 0.08 | -0.2 ± 0.3 |
| Copepods | -0.25 ± 0.29 | -0.02 ± 0.03 | -0.08 ± 0.07 |
| ANOVA Results: Treatment: $F_{(2,27)} = 0.15$; $p = 0.86$, $n = 12$<br>Zooplankton Taxa: $F_{(2,27)} = 1.96$; $p = 0.17$, $n = 12$<br>Treatment:Zooplankton: Taxa: $F_{(4,27)} = 0.48$; $p = 0.75$, $n = 12$ | | | |

## Discussion

The transition from endogenous to exogenous feeding is a ‘critical period’ in larval fish development, characterized by high mortality rates that influence longer-term population growth (Fuiman and Werner 2002). Here, we demonstrate that browning may add stress to this critical stage by decreasing zooplankton prey availability. Browning did not, however, directly alter larval fish feeding efficiency or prey selectivity, suggesting plasticity in foraging behavior under varying light conditions. To our knowledge, this is the first study examining the effects of freshwater browning on larval fishes. Understanding the balance between direct versus indirect effects of browning on early life history stages will improve fish conservation and management strategies in response to continued organic matter loading with changes in climate and land use.

### Indirect Effects of Browning

The transfer of energy through the traditional grazer food chain depends on basal resources to support higher trophic levels. Therefore, one of the most profound effects of freshwater browning is the reduction in primary production, due to increased light attenuation (Wetzel 2001; Thrane et al. 2014; Solomon et al. 2015), that subsequently reduces energy flow to zooplankton and fish (Jones et al. 2012; Solomon et al. 2015; Creed et al. 2018). Our study supports these general observations and raises concerns for larval fish. With increasing brown color, we observed a reduction in the quantity of light, decreased phytoplankton biomass, and decreased zooplankton densities. In turn, the foraging rate, growth, and survival of larval largemouth bass declined.

Starvation affects larval fish more than juvenile or adult stages because of their high metabolic demands coupled with low energy reserves in their tissues (Fuiman 2002). Within 4 - 5 days at 25 - 30 °C, larvae will likely starve to death given low to no food rations (Fuiman 2002). In the present study, after 6 days at 22 - 23°C, fish larvae in the moderate and dark brown treatments exhibited a 33% mortality rate while the surviving larvae in these treatments displayed concave-shaped, empty stomachs. Although we did not quantify gut fullness, these observations suggest a positive relationship between browning and starvation risk. Fish larvae were found dead in the darker brown treatments near the end of the experiment (Day 5 or 6) when zooplankton densities were already visibly depleted.

At the time fish larvae were introduced to our experimental tanks, zooplankton densities in the moderate and dark brown treatments were 2 to 4 times less than in the light brown and lake water treatments (i.e. 46 - 72 individuals per liter compared to 120 - 190 individuals per liter). Daily consumption rates for larval fish are often greater than a larvae’s own biomass (Post 1990). We did not directly measure zooplankton or larval fish biomass in our study. However, based on published length-weight regressions (larval fish: Brecker 1993; zooplankton: reviewed in Watkins et al. 2011), we roughly estimate that larval fish biomass was ^~^2020 *μg* dry weight per fish at the time of their introduction to the experimental tanks while total zooplankton biomass was ^~^1577 ± 447 *μg* dry weight in the dark brown treatment, ^~^2965 ± 361 *μg* dry weight in the moderate brown treatment, ^~^6300 ± 1582 *μg* dry weight in the light brown treatment, and ^~^4962 ± 228 *μg* dry weight in the lake water treatment. This resulted in a ratio of zooplankton:fish biomass of 0.8, 1.5, 3.1, and 2.5 in the dark brown, moderate brown, light brown, and lake water treatments, respectively. After 24 hours, fish larvae consumed approximately 0.26, 0.54, 1.0, and 1.5 grams of zooplankton per gram of fish in the dark brown, moderate brown, light brown, and lake water treatments, respectively. Based on data from Houde and Zastrow (1990), freshwater fish should consume a minimum of ^~^0.5 - 0.7 grams of prey per gram of fish per day at 23 °C to meet mean weight-specific growth rates. This suggests that larvae in the moderate and dark brown treatments were food limited.

We placed two larvae in each tank to encourage routine behavior, which resulted in a fish density of ^~^118 larvae m^−3^. This is relatively high but not uncommon in nature, particularly during the early months of fish hatching (Santucci et al. 2003). Previous studies have shown that when zooplankton consumption outweighs zooplankton reproduction, fish larvae may rapidly deplete their food source, affecting future growth and survival (Mehner and Thiel 1999; Santucci et al. 2003; Hansson et al. 2007). This may occur more often in brown systems if zooplankton abundances are low at the time of fish hatching. Furthermore, as surface water temperatures rise as a result of global climate change, fish feeding rates are predicted to increase as a consequence of higher metabolic rates (van Dorst et al. 2019). Brown waters are likely to increase in temperature more rapidly due to the increased absorption of solar radiation (Solomon et al. 2015), further exacerbating competition for limited resources in these systems.

Unlike prey abundance, we did not see significant differences in zooplankton community composition with increased water color. Others studies have also noted minimal changes in zooplankton community composition with freshwater browning, and there does not seem to be a clear pattern in positive versus negative effects on specific zooplankton groups (Nicolle et al. 2012; Ekvall and Hansson 2012; Kelly et al. 2014; Robidoux et al. 2015; Lebret et al. 2018; Leech et al. 2018). In the present study, the zooplankton community shifted from copepod to cladoceran dominated across all treatments, with smaller-bodied *Bosmina* the dominant zooplankton prior to larval fish introduction. Abundances of large-bodied *Daphnia* were similar in all treatments during the first two weeks of the experiment, but for unknown reasons, continued to increase only in the light brown treatment (i.e., ^~^ 30 individuals per L at Day 28). It is possible that the light brown treatment provided the ideal diet for *Daphnia*, with sufficient carbon and nutrients from both algal- and terrestrially-derived resources (Lennon et al. 2013; Gall et al. 2017; Tang et al. 2018).

Overall, the zooplankton community that developed in our experiment provided a standard diet for fish larvae. Typically, larger crustacean zooplankton promote foraging success, growth, and survival (Crowder et al. 1987; Mills et al. 1989). However, for younger larvae that are gape-limited, smaller prey, like *Bosmina* spp., can be more beneficial (Mehner and Thiel 1999). Nevertheless, zooplankton prey abundance in the moderate and dark brown treatments was too low to support larval development.

Similar to previous studies (Batt et al. 2015; Karlsson et al. 2015; Solomon et al. 2015), reduced zooplankton abundance coincided with reduced phytoplankton biomass as browning increased. Interestingly, despite having similar water color, the light brown and lake water treatments displayed opposite trends in phytoplankton biomass after Day 14. In the lake water treatment, increased zooplankton abundance likely resulted in greater consumption of phytoplankton biomass. However, we are uncertain why the same pattern did not occur in the light brown treatment. It is possible that the light brown treatment, made with the COMBO medium, provided phytoplankton with more nutrients for growth to keep up with zooplankton consumption. We were unable to measure nutrient concentrations at the time of these experiments; however, water quality monitoring data collected by the Virginia Department of Environmental Quality for Sandy River Reservoir indicate generally lower nitrogen and phosphorus concentrations than present in the COMBO medium.

Phytoplankton species composition and nutritional quality were not assessed as part of our study; however, we recognize that these factors can also influence energy and nutrient availability for zooplankton, and consequently larval fish (Taipale et al. 2018; Creed et al. 2018). Because of logistics, we chose to use an artificial assemblage of green algal species that are generally favorable foods. In nature, phytoplankton communities can shift towards cyanobacteria with increased water color, which could further reduce zooplankton densities (Ekvall et al. 2013; Robidoux et al. 2015). It is also possible that the algae used in our experiment may not have been adapted to the low light conditions of brown water systems given that they were purchased from Carolina Biological. However, our results are comparable to field studies examining natural communities along a water color gradient (Ask et al. 2009; Karlsson et al. 2009; Thrane et al. 2014).

Reductions in prey consumption, and possibly less nutritious prey, ultimately led to reductions in fish growth. Fish larvae in the lake water and light brown treatments, on average, grew twice as fast as the larvae in the moderate brown treatment and four times faster than the larvae in the dark brown treatment. Although, we recognize that there was relatively high unexplained variability in growth within our treatments (e.g. fish in two replicate tanks of the moderate brown treatment exhibited no growth while the third replicate exhibited relatively high growth). Growth rates are important for several reasons. Larval fish become better at avoiding predators and detecting prey as swimming strength and visual acuity increases with body size (Fuiman 2002). In addition, larger larvae consume bigger, more energy-rich prey. Growth in juvenile and adult fish is often slower in brown compared to blue lakes, resulting in a smaller length-at-age (Estlander et al. 2010; Horppila et al. 2010; Benoit et al. 2016; van Dorst et al. 2018), and our data suggest similar patterns for larval fish. These reductions in individual growth lead to further reductions in population growth and fish biomass as lakes darken in color (Karlsson et al. 2009, 2015; Finstad et al. 2014), which can alter food web structure and lower fisheries yields.

Importantly, the results of our indirect effects experiment were influenced by the timing of fish introduction at Day 28. If the fish had been introduced at earlier time points, when zooplankton densities were similar in all treatments, we likely would have observed no effect of browning on larval fish. Indeed, in systems with low prey availability, fish foraging, growth, and survival are reduced regardless of water color (Mehner and Thiel 1999; Santucci et al. 2003; Hansson et al. 2007). Nonetheless, similar to our experimental results, browning has been demonstrated in field and laboratory studies to reduce zooplankton densities (e.g. Ekvall et al. 2013; Robidoux et al. 2015; Leech et al. 2018). Moreover, it is often stated that these differences in prey availability affect fish growth and survival, but experimental evidence is limited. Here, we provide quantitative data on how alterations in prey densities, due to increased browning, may affect larval fish foraging, growth, and survival in nature. We argue that these valuable data can served as a starting point to design larger-scaled mesocosm experiments or observational studies.

### Direct Effects of Browning

When given an equal abundance of zooplankton prey, neither the foraging efficiency nor prey selectivity of larval largemouth bass or bluegill were affected by increased browning. Our results are similar to Stasko et al. (2012), which found no significant effect of browning on juvenile roach feeding rates but are contrary to Jönsson et al. (2013), which reported decreases in reactive distance and capture success of piscivorous Northern pike (*Esox lucius*) feeding on roach (*Rutilus rutilus*) with increased water color. Weidel et al. (2017) reported significant effects of browning on juvenile largemouth bass and bluegill foraging, but water color explained only ^~^25-28% of the variation in foraging rates. Combined, results from our study and others (Estlander et al. 2012; Ranaker et al. 2012; Nurminen et al. 2014) suggest that effects of browning on fish foraging rates may be age- or species-specific.

Light intensities across all our treatments (i.e. ^~^15 – 35 μmol m^−2^ s^−1^ PAR) may have been adequate for larval fish foraging, resulting in the lack of direct effects. For example, Miner and Stein (1993) reported that larval bluegill successfully feed on zooplankton at light intensities above 450 lux (i.e., ^~^8 μmol m^−2^ s^−1^ based on the conversion factor for sunlight). Some have suggested that reductions in light levels have a greater effect on later life history stages of fish because of the positive relationship between sighting distance and body size (Askne and Giske 1993; Fiksen et al. 2002). If browning does directly affect visual foraging, it may do so by limiting the thickness and daily duration of the photic zone, such that there is a spatiotemporal contraction of foraging habitat critical to growth and survival. Additionally, the metabolic costs associated with searching for prey may increase with reductions in light levels, particularly if prey availability declines with increased browning.

Larval fish used in our experiments may have been pre-adapted to the low-light environment of brown water systems. In general, the man-made reservoirs and ponds of central Virginia, USA have relatively high light attenuation due to increased cDOM inputs and/or increased phytoplankton growth (Figure S1). Compared to local lakes and ponds in the region, our dark brown treatment increased water color approximately 10 times. However, we may have seen direct effects of browning on fish feeding had we continued to add organic matter to the tanks. Moreover, fish inhabiting clear, blue lakes may show a greater response to freshwater browning than those in our study (Stasko et al. 2015), and we encourage further experimentation with larvae from these systems.

Because foraging success plays a critical role in survival and reproduction, there is likely strong selection pressure to adapt to changing environmental conditions. As light availability for visual foraging declines with browning, fish may shift from vision to other sensory mechanisms, such as mechanoreception or olfactory cues, to detect zooplankton prey. For example, previous research ablating the function of superficial neuromasts with neomycin or streptomycin has shown reduced feeding rates in marine fish larvae (Jones and Janssen 1992; Cobcroft and Pankhurst, 2003; Sampson et al. 2013). Larval zebrafish have been shown to learn to use mechanoreception to feed in the dark (Carillo and McHenry 2016). There is also recent evidence that juvenile bluegill may actually feed more in the open pelagic water column at night and horizontally migrate towards the littoral zone during the day to avoid piscivorous predators (Shoup et al. 2014). Future research should investigate the potential for alternative feeding strategies as waters brown in color and be cautious about our biases as human researchers relying on light and vision (Cumming et al. 2018).

### Small Enclosures versus Natural Systems

We recognize that our experiment was conducted in relatively small containers and that caution must be applied when scaling up results to natural systems. Nevertheless, our observations of phytoplankton, zooplankton, and fish responses to browning are similar to those reported by recent field-based studies (Karlsson et al. 2009, 2015; Finstad et al. 2014 and others mentioned above), providing confidence in our results. Working with fragile fish larvae is challenging, and smaller containers minimize some of the logistical constraints. Yet, the small, confined tank could have influenced feeding rates. Given the size of larvae used in our experiments (10 - 13 mm), their visual acuity, or reaction distance, is approximately one body length (Werner 1969). However, we do not know how long it took larvae to search the tank. Larger containers may have revealed greater differences in feeding rates across treatments, similar to Weidel et al. (2017).

We also did not quantify the spatial distribution of zooplankton in the tanks, which could have influenced feeding rates. Experimental tanks were covered during the feeding trials to avoid distracting the fish. Zooplankton are known to swim downward in the presence of light and fish, and it is possible that the zooplankton could have clumped at the bottom of the tanks, making them easier to find and capture. However, this behavioral pattern in the zooplankton was not obviously apparent to us while breaking down the tanks.

Interestingly, Seekell et al. (2018) reported that the relationship between fish biomass and water color was more negative in deep lakes compared to shallow lakes within the boreal region of Sweden. The authors state that the negative effects of browning associated with light extinction and decreased primary production are minimized in shallow lakes because light often reaches the lake bottom. Moreover, most fish inhabiting deeper lakes were observed in the littoral zone. Moving into shallow waters may provide fish adequate light to forage in brown waters, as supported by our results in shallow, experimental tanks (i.e., ^~^0.2 m depth).

### Implications and Applications

Given increasing trends in temperature and precipitation with global climate change, the browning of inland and coastal waters is predicted to continue, with far reaching consequences for aquatic ecosystems (de Wit et al. 2016; Weyhenmeyer et al. 2016). Here, we demonstrate that freshwater browning reduces energy flow to higher trophic levels, negatively affecting larval fish growth and survival during a ‘critical period’. Because of their ecological and economic value, fish population dynamics are closely monitored and predicted by natural resource managers and conservationists. While current population growth models focus on the importance of temperature and nutrients (e.g., Deslauriers et al. 2017), our study and others, highlight the need to incorporate water color (i.e. cDOM) to more accurately predict recruitment strength, sustainable yields, and food web stability in systems affected by browning.

## Acknowledgements

We thank David and Navona Hart for granting access to their pond and Mark Rolfing for his assistance in counting bacteria samples. Funding for this research came from the Longwood University PRISM Summer Research Program and a faculty development grant to DML from Longwood University. Collection of larval bluegill and largemouth bass was approved by the Virginia Department of Game and Inland Fisheries, Permit# 051397. This manuscript was improved by thoughtful comments from Sönke Johnsen, Tessa DeWalt, and two anonymous reviewers.

## Supplementary Materials

**Figure S1.**
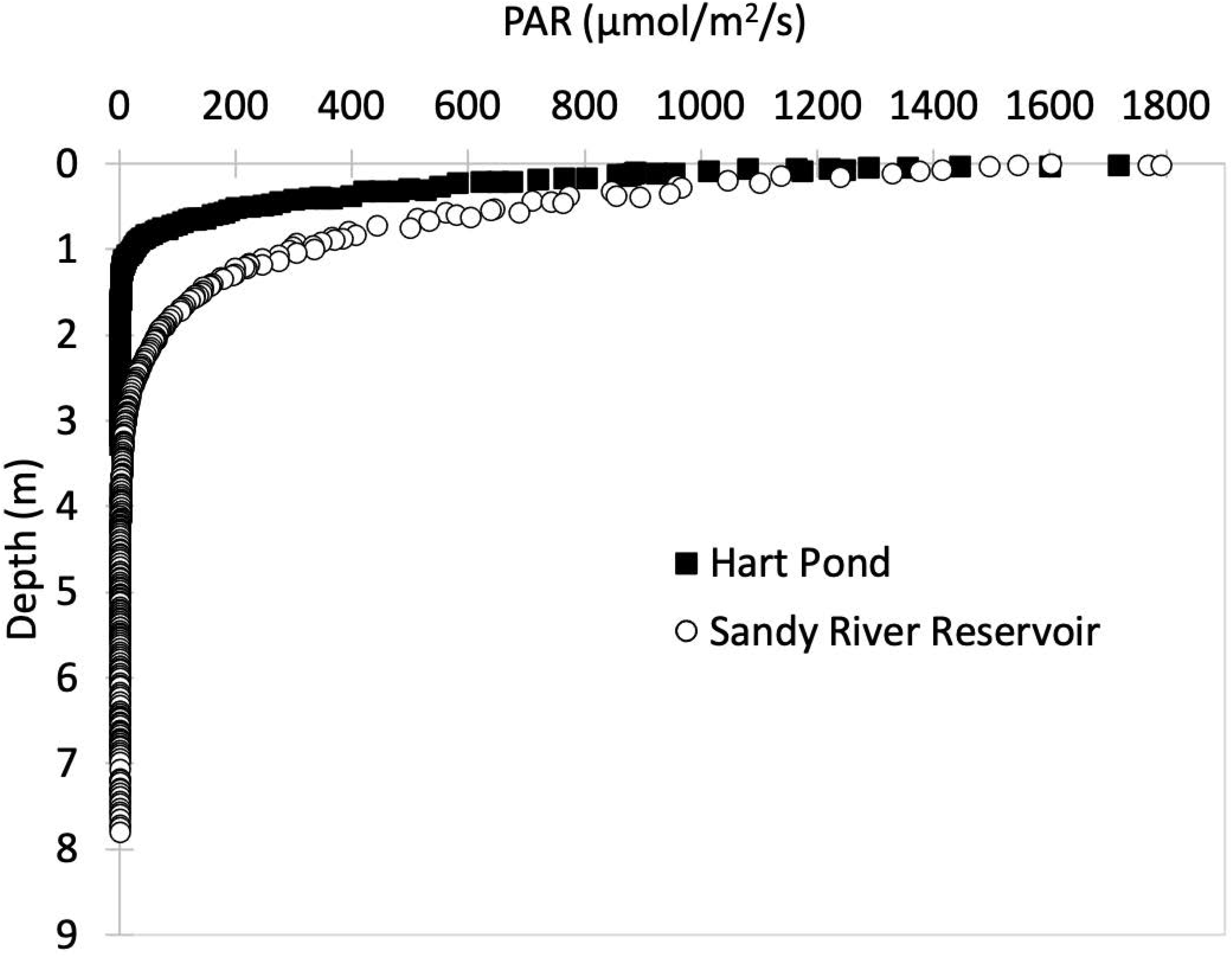
Representative downwelling irradiance curves for photosynthetically active radiation (PAR) in Sandy River Reservoir and Hart Pond. Both are located near Farmville, VA, USA and were used as our study systems to collect larval fish and zooplankton. Measurements were taken with a Biospherical Instruments, Inc. BIC profiling radiometer in June 2014.

**Figure S2.**
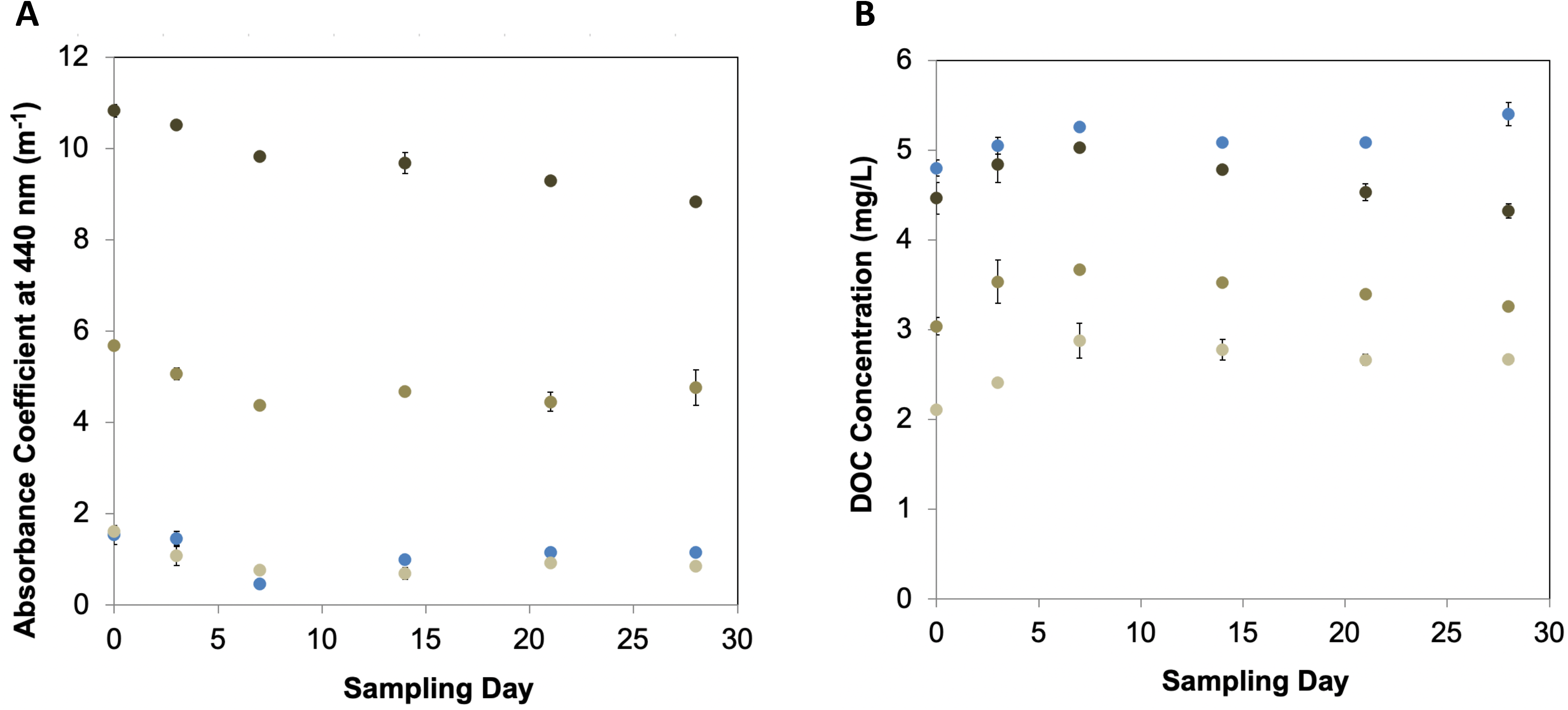
Mean ± S.E. of (A) a_440_ and (B) dissolved organic carbon concentrations over time during the Indirect Effects experiment. Data were collected at Day 0, 3, 7, 14, 21, and 28 as described in the Methods.

